# The three dimensional structure of Bovine Salivary Protein 30b (BSP30b) and its interaction with specific rumen bacteria

**DOI:** 10.1101/448167

**Authors:** Heng Zhang, Judith Burrows, Graeme L. Card, Graeme Attwood, Thomas T. Wheeler, Vickery L. Arcus

**Author notes:** Current address: 35 Prisk Street, Melville, Hamilton 3206, New Zealand (Graeme L. Card).

## Abstract

Bovine Salivary Protein 30b (BSP30b) is a member of the tubular lipid-binding (TULIP) superfamily that includes the human bactericidal/permeability-increasing proteins (BPI), lipopolysaccharide binding proteins (LBP) and palate, lung, and nasal epithelium carcinoma-21 associated proteins (PLUNC). BSP30b is most closely related to the PLUNC family and is predominantly found in bovine saliva. There are four BSP30 isoforms (BSP30a-d) and collectively, they are the most abundant protein component of bovine saliva. The PLUNC family members are proposed to be lipid binding proteins, although in most cases their lipid ligands are unknown. Here, we present the X-ray crystal structure of BSP30b at 2.0 Å resolution. We used a double methionine mutant and Se-Met SAD phasing to solve the structure. The structure adopts a curved cylindrical form with a hydrophobic channel formed by an α/β wrap, which is consistent with the TULIP superfamily. The structure of BSP30b in complex with oleic acid is also presented where the ligand is accommodated within the hydrophobic channel. The electron density for oleic acid suggests that the ligand is only partially occupied in the binding site implying that oleic acid may not be the preferred ligand. GFP-tagged BSP30b binds to the surface of olive oil droplets, as observed under fluorescent microscopy, and acts as a surfactant consistent with its association with decreased susceptibility to bloat in cattle. Bacteria extracted directly from bovine rumen contents indicate that the GFP_BSP30b fusion protein binds to a small number of selected bacterial species *in vivo*. These results suggest that BSP30b may bind to bacterial lipids from specific species and that this abundant protein may have important biological roles via interacting with rumen bacteria during feeding and rumination.

## Introduction

PLUNC (<u>p</u>alate, <u>l</u>ung, and <u>n</u>asal epithelium <u>c</u>arcinoma-associated) proteins are a family of proteins predominantly expressed in the upper respiratory tract, nasal mucosa, and oral cavity in mammals. PLUNC proteins are further subdivided into LPLUNC (long PLUNC) proteins and SPLUNC (short PLUNC) proteins (1). Although several studies have characterized the expression of PLUNC proteins, their biological function(s) are not well understood. Studies have reported that PLUNCs can bind to bacteria and LPS (lipopolysaccharide) (2-4) but in antimicrobial assays PLUNCs showed no bactericidal activity against *Escherichia coli DH5-a, P. aeruginosa PA01* or *Listeria monocytogenes* (5).

From a structural point of view, the PLUNC protein family belongs to the TULIP (tubular lipid-55 binding) superfamily (6). The tertiary structure of this domain is a long, bent helix wrapped in four highly curved anti-parallel β-strands, forming a central hydrophobic channel ideal for lipid binding (7). The TULIP superfamily consists of three protein families – BPI-like, SMP-like (synaptotagmin-like, mitochondrial and lipid-binding protein), and Takeout-like protein families. The BPI-like family includes BPI, LBP, CETP (the cholesteryl ester transfer protein), PLTP (the phospholipid transfer protein), and PLUNC. These proteins are either involved in innate immunity against bacteria through their ability to bind LPS, or in lipid exchange between lipoprotein particles. The Takeout-like protein family consists of various arthropod allergens and insect juvenile hormone-binding proteins, which transport lipid hormones to target tissues during insect development. The SMP-like protein family includes subunits of the ERMES complex (ER-mitochondria encounter structures) and the extended synaptotagmins (E-Syts), which appear to be mainly located at membrane contact sites between organelles, mediating inter-organelle lipid exchange (7).

The first described PLUNC protein in cattle was named Bovine Salivary Protein 30 (BSP30), based on its mass of approximately 30 kDa (8). This “BSP30” protein is one of the most abundant proteins in bovine saliva, with a concentration of 0.1-0.5 mg.ml^-1^. Its abundance was proposed to be associated with the susceptibility of cattle to pasture bloat, a metabolic disease characterized by build-up of stable foam in the rumen and hence impairment of the eructation mechanism, causing rumen distension and respiratory distress (9). Although BSP30b has been systematically renamed as BPIFA2B in 2011 (10), in this paper, BSP30b is used for the sake of simplicity and to remain consistent with previous publications on this particular protein.

Further investigation of BSP30 revealed that it is a mixture of two closely related proteins - 78 BSP30a and BSP30b, sharing 83% amino acid identity, with expression restricted to salivary tissue (11). Sequencing analysis of the PLUNC gene locus within cattle revealed that there are thirteen genes within this locus, nine of which are orthologous to genes in the human and mouse PLUNC gene locus (12). The other four genes, namely *BSP30a*, *BSP30b*, *BSP30c*, and *BSP30d*, are thought to arise through gene duplications and are a characteristic feature of ruminants. The implication is that these proteins contribute to ruminant-specific physiological functions (12, 13). A gene expression study indicated that the mRNAs of these four *BSP30* genes are most abundant in tissues associated with the oral cavity and airways. Notably, *BSP30a* and *BSP30b* are abundantly expressed in the salivary gland independently of one another and the proteins are secreted into saliva, accounting for 15-30% of the total protein in bovine saliva (14).

Analyses of the cattle PLUNC family at the protein level have focused on BSP30a and BSP30b. Immunohistochemical analysis of bovine salivary glands with antibodies specific for both BSP30a and BSP30b suggested that they are localized to the serous secretory cells (15). Further analysis with Western blotting revealed that BSP30b is present in the parotid, submandibular and buccal salivary glands; whilst BSP30a is only present in the submandibular gland, although its mRNA was shown to be expressed abundantly in the parotid (13, 14). Functional studies suggests that both BSP30a and BSP30b can suppress the growth of *Pseudomonas aeruginosa* with moderate potency but suppression was not observed for other pathogens tested, indicating that, unlike the BPI-family proteins, the BSP30 proteins may not have a primary function as bactericidal proteins (13, 15). It has also been observed that both BSP30a and BSP30b have no significant LPS binding activity, suggesting the mechanism of their moderate antibacterial activity is independent of LPS opsonisation (13).

To address functional questions regarding BSP30 proteins, we have solved the X-ray crystal structure of BSP30b at 2.0 Å resolution. The structure shows the conserved architecture seen among the TULIP superfamily. We have also solved the structure of BSP30b in complex with oleic acid which shows that the hydrophobic channel can sequester one molecule after co-105 crystallization. However, the electron density for oleic acid is relatively poor, suggesting it is not the preferred lipid ligand of BSP30b. We demonstrate that olive oil can be dispersed into nano-droplets after mixing with BSP30b and thus we corroborate its surfactant properties. We further report that BSP30b binds to the surface of a small subset of rumen bacterial species and thus, BSP30b may bind to a specific range of lipid ligands and play a biological role in the rumen via interaction with specific rumen bacteria.

## Materials and methods

### Protein expression, crystallization, and structure determination

The double mutant Bovine BSP30b gene containing two methionine residues (replacing phenylalanine residues, *BSP30b_F52M_F106M*) was synthesized with restriction sites BamHI and XhoI by GeneArt (Life Technologies, Carslbad CA). This gene was ligated into the BamHI/XhoI site of the *pProEx Htb* vector and transformed into *Escherichia coli BL21* cells. Seleno-methionine (SeMet) incorporated BSP30b_F52M_F106M was expressed in M9 minimal media. A cell culture was grown at 37 °C with 100 μg.mL^-1^ ampicillin with shaking at 200 rpm. When the OD_600_ of the culture reached 0.4-0.6, 100 mg of lysine, phenylalanine and threonine, 50 mg of isoleucine and valine, and 60 mg of selenomethionine were added and the growth was continued for a further 15 min. IPTG (0.5 mM final concentration) was added for induction of protein expression and the culture was grown overnight. Cells were harvested by centrifugation at 6000 rpm for 20 min and resuspended in 20 ml of lysis buffer (20 mM HEPES pH 8, 150 mM NaCl, 1 mM β-mercaptoethanol, 20 mM imidazole) and then sonicated on ice. Disrupted cells were centrifuged at 13,000 rpm and the soluble fraction containing BSP30b_F52M_F106M SeMet was loaded onto a HiTrap Ni column for protein purification. Protein was further purified with anion exchange chromatography using MONO Q 4.6/100 PE (GE Healthcare). The protein was finally purified with size exclusion chromatography (SEC) using a Superdex 75 16/60 column (GE Healthcare) and eluted with 10 mM HEPES pH 8, 150 mM NaCl. The purified protein was concentrated to 8 mg.mL^-1^ for crystallization.

The BSP30b_F52M_F106M SeMet derivative was crystallized using hanging drop vapour diffusion at 18 °C by mixing 1.5 μL of the protein solution with 1.5 μL of the reservoir solution (0.1 M HEPES pH 7.5, 0.2 M CaCl_2_, 9 % PEG 3350, and 5 % isopropyl alcohol). Crystals were obtained after two weeks and were flash frozen in liquid nitrogen without cryo-protectant for data collection. Single-wavelength anomalous dispersion (SAD) data were collected using X-rays at a wavelength of 0.9537 Å at the Australian Synchrotron MX2 beamline with an ADSC Quantum 210r detector. Data were integrated using iMosfilm (16) and scaled using Aimless (17) within the CCP4 platform suite 6.4.0 (18). Phases were solved by using AutoSol (19) and the initial structure was built using AutoBuild (20) within the PHENIX platform suite (21). The structure was first refined using *phenix.refine* (22) in PHENIX. Subsequent cycles of manual refinement using COOT 7.0 (23) and automatic refinement in PHENIX were performed.

### Co-crystallization of BSP30b_F52M_F106M with oleic acid

To obtain crystals of an BSP30b_F52M_F106M-oleic acid complex, purified protein was concentrated to ∼2 mg.mL^-1^ and incubated with 20 μL of oleic acid at 4 °C overnight. This mixture was purified using SEC to remove unbound oleic acid and then concentrated to 8 mg.mL^-1^ for crystallization. Crystals were obtained after two weeks under the same conditions as for the protein alone. A complete diffraction dataset was collected at the Australian Synchrotron MX2 beamline using the new ACRF Eiger 16M detector (24). The structure was solved by molecular replacement using Phaser (25) with the solved BSP30b structure as the reference model. The structure was built and refined as above.

### Fluorescent microscopy of GFP_BSP30b mixed with olive oil

A GFP_BSP30b fusion protein was constructed to visualize the interaction between BSP30b and olive oil, as well as the interaction with microorganisms in the rumen. Both GFP and BSP30b were cloned into *pProEx Htb* with the BamHI/PstI and XhoI/HindIII restriction sites, respectively. The fused protein was expressed in LB media with a 6 x histidine tag at the N-158 terminus.

Since the primary lipid source for cattle is long chain fatty acids in triglyceride form, the interaction between BSP30b and triglyceride was studied to illustrate the function of BSP30b. Olive oil was chosen for this purpose as it is composed mainly of the mixed triglyceride esters of oleic acid, linoleic acid, palmitic acid and other fatty acids, similar to the composition of triglycerides from pasture grass.

20 μL of olive oil was added to 500 μL of GFP_BSP30b at a concentration of 2 mg.mL^-1^, which was then vortexed for 2 mins and left on the lab bench for 20 mins for equilibration. 10 μL of this mixture was pipetted onto a glass slide and then covered with a cover slip. The slide was viewed by using a ZEISS AXIOSTAR (LLC, US) bench top microscope. Images were captured at 40 X magnification with filter 1 (blue light) using a Nikon Coolpix 4500 camera (Tokyo, Japan).

### Fluorescent microscopy of GFP_BSP30b with rumen samples

Rumen fluid was collected from a fistulated cow at AgResearch (Palmerston North, New Zealand). After filtration through cheesecloth to remove large feed particles, the fluid was incubated at 39 °C in a waterbath and kept anaerobic by bubbling with a slow stram of CO_2_. This filtered fluid was further separated into bacterial and protozoal fractions by low speed centrifugation at 300 x g for 5 min.. The pellet, which contained the protozoa and some bacteria, was diluted with Artificial Saliva (142 mM Na_2_HPO_4_, 84 mM NaHCO_3_, 100.1 mM KHCO_3_, 60.1 mM Urea, and 58.4 mM NaCl) to the same volume as the supernatant. The supernatant, which then contained the bulk of the bacteria, was centrifuged at 15,000 x g for 15 min to pellet the bacteria. The bacterial pellet was then re-suspended with Artificial Saliva to the same volume as the supernatant. BSP30b protein (0.1 mg.mL^-1^) was then mixed with the protozoal or bacterial fractions at a ratio of 1:1 by volume. 20 μl of each mixture was pipetted on a slide, covered with a coverslip and viewed under ultraviolet (UV) illumination using a Leica Model DN 2500 microscope at 100x magnification under oil..

## Results

### Structure determination and description

Crystals of a double Phe-to-Met mutant of BSP30b diffracted to 2.0 Å resolution with a space group of C2 and unit cell dimensions: a = 81.65 Å, b = 59.57 Å, c = 89.94 Å, α = ν = 90°, β = 106.25° (Table 1). The structure was solved using SeMet incorporated protein and SAD methods.

**Table 1.** Crystallographic data collection and refinement statistics.

|  | BSP30b_F52M_F106M | BSP30b_F52M_F106M-oleic acid |
| --- | --- | --- |
| **Data collection** |  |  |
| Wavelength | 0.9537 Å | 0.95369 Å |
| Space group | C121 | C121 |
| Cell constants<br>a, b, c, $\alpha$, $\beta$, $\gamma$ | 81.65Å 59.57Å 89.94Å<br>90.00° 106.25° 90.00° | 83.75Å 61.40Å 91.09Å<br>90.00° 107.74° 90.00° |
| Resolution (Å) | 47.43 – 2.00 (2.11 – 2.00) | 86.76 – 1.97 (2.02 – 1.97) |
| $R_{merge}$ | 0.100 (0.622) | 0.067 (0.683) |
| Completeness (%) | 99.9 (100) | 99.7 (99.7) |
| Redundancy | 7.2 (7.3) | 5.7 (6.1) |
| No. of observations | 201779 (29708) | 177824 (13279) |
| No. of unique reflections | 28136 (4080) | 31005 (2186) |
| Mean $I/\sigma(I)$ | 11.8 (3.5) | 9.9 (2.3) |
| Anomalous completeness (%) | 98.5 (98.1) |  |
| Anomalous multiplicity | 3.7(3.8) |  |
| **Refinement** |  |  |
| $R_{work}$ | 0.2051 | 0.2326 |
| $R_{free}$ | 0.2514 | 0.2878 |
| Average B, all atoms (ligand) (Å<sup>2</sup>) | 36.0 | 55.9 (63.5) |
| Ramachandran favoured (outliers) | 97.05% (0.00%) | 94.29% (0.54%) |
| Rotamer outliers | 0.00% | 0.32% |

BSP30b crystallized with two molecules in the asymmetric unit (Fig. 1a). Each molecule consists of 7 α helices and 6 β strands as well as connecting turns and loops, which constitute the conserved TULIP fold (Fig. 1b). The N-terminus of the structure of BSP30b contains four consecutive helices (α1, residues 40-48; α2, residues 50-53; α3, residues 55-69; α4 residues 81-100) with the fourth helix lying antiparallel to the preceding three (Fig. 1c). Electron density for the loop joining α3 and α4 is absent. The central section of the structure comprises a highly twisted 4-stranded antiparallel β sheet where the first and fourth strands are interrupted by a loop and a short helix respectively (Fig. 1c). A long curved α helix (α6 residues 193-224) follows the central section and the structure terminates with a C-terminal helix unique to the BSP30 subfamily (α7 residues 227-235).

**Figure 1.**
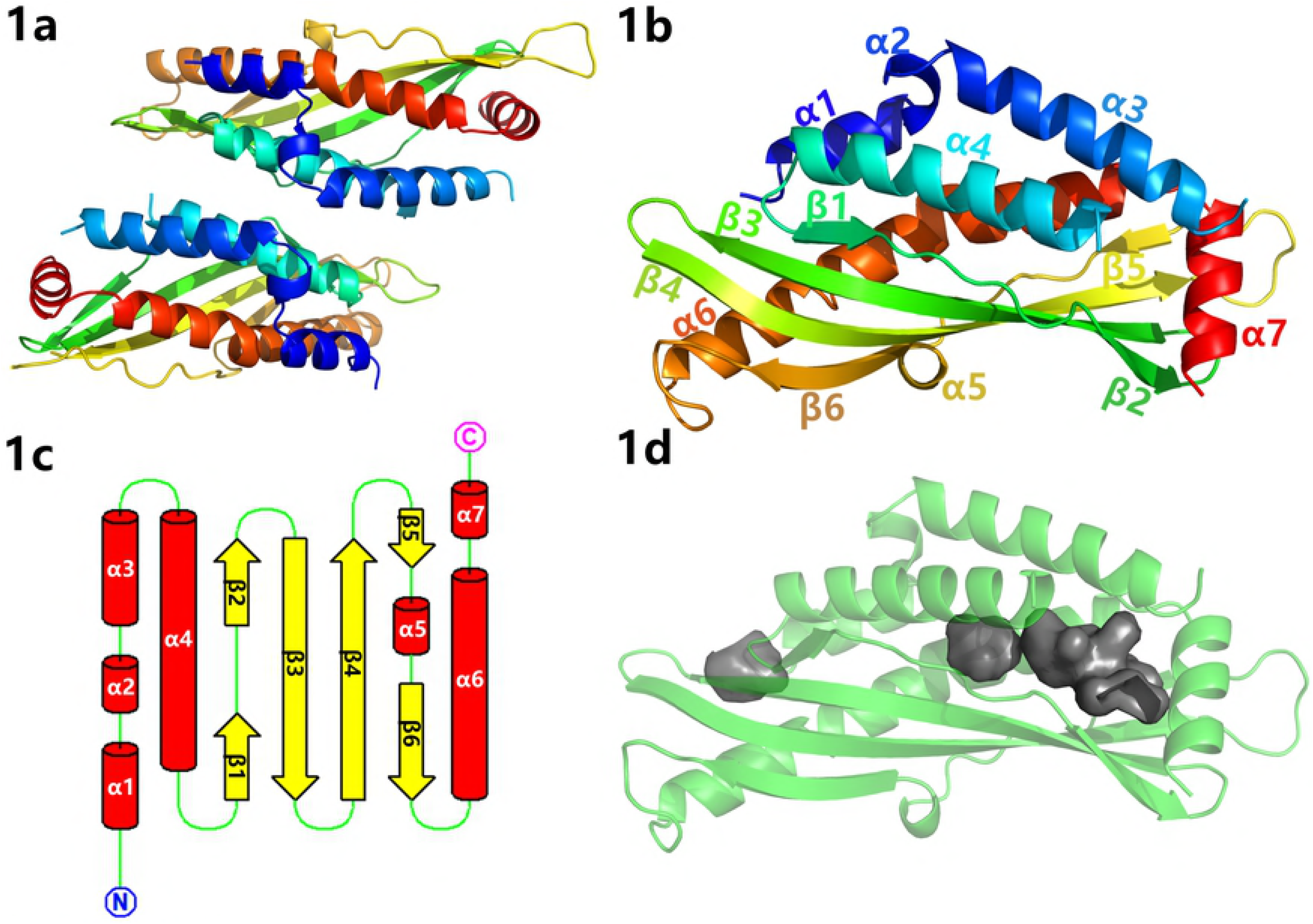
Structure of BSP30b. 1a, asymmetric unit composed of two copies of BSP30b molecule. 1b, the tertiary structure of apo-BSP30b, coloured in rainbow mode ranging from blue (N terminus) to red (C terminus). 1c, topology diagram of BSP30b. All seven E helices are represented by red cylinders; six β strands by yellow arrows; green lines represent the loops and turns. 1d, representation of the internal cavity, calculated using PyMOL (Delano Scientific).

The hydrophobic channel of BSP30b (∼22.6 Å in length) is enclosed by all of the secondary structure elements. This internal cavity has an opening located adjacent to the C-terminal helix at one end of the long axis of the structure (Fig 1d). The flexible loop between helices α3 and α4 at the opening of the channel potentially caps the channel and may play a role in ligand binding. BSP30b also has a conserved disulfide bridge common to the PLUNC family (cysteines 172 and 225) that joins the long curved helix α6 and the loop region between helix α5 and strand β5. This disulfide bond has been proposed to contribute to the stability of the overall structure (26).

### Structural comparison

The BSP30b structure was compared with other structures in the PDB using PDBeFold. From the structural alignment, two proteins are related: SPLUNC1 from human (Fig. 2a, PDB code 4KGH) with a Z score of 4.8 and a root mean squared deviation (RMSD) for Cα atoms of 3.2 Å (a total of 144 residues aligned); and BPIFA1 from mouse with a Z score of 5.4 and RMSD for Cα positions of 3.4 Å (total of 151 residues aligned). All three proteins have the archetypal tubular structure with the conserved hydrophobic channel and the conserved disulfide bond (Fig. 2a and 2b) at the same position.

**Figure 2.**
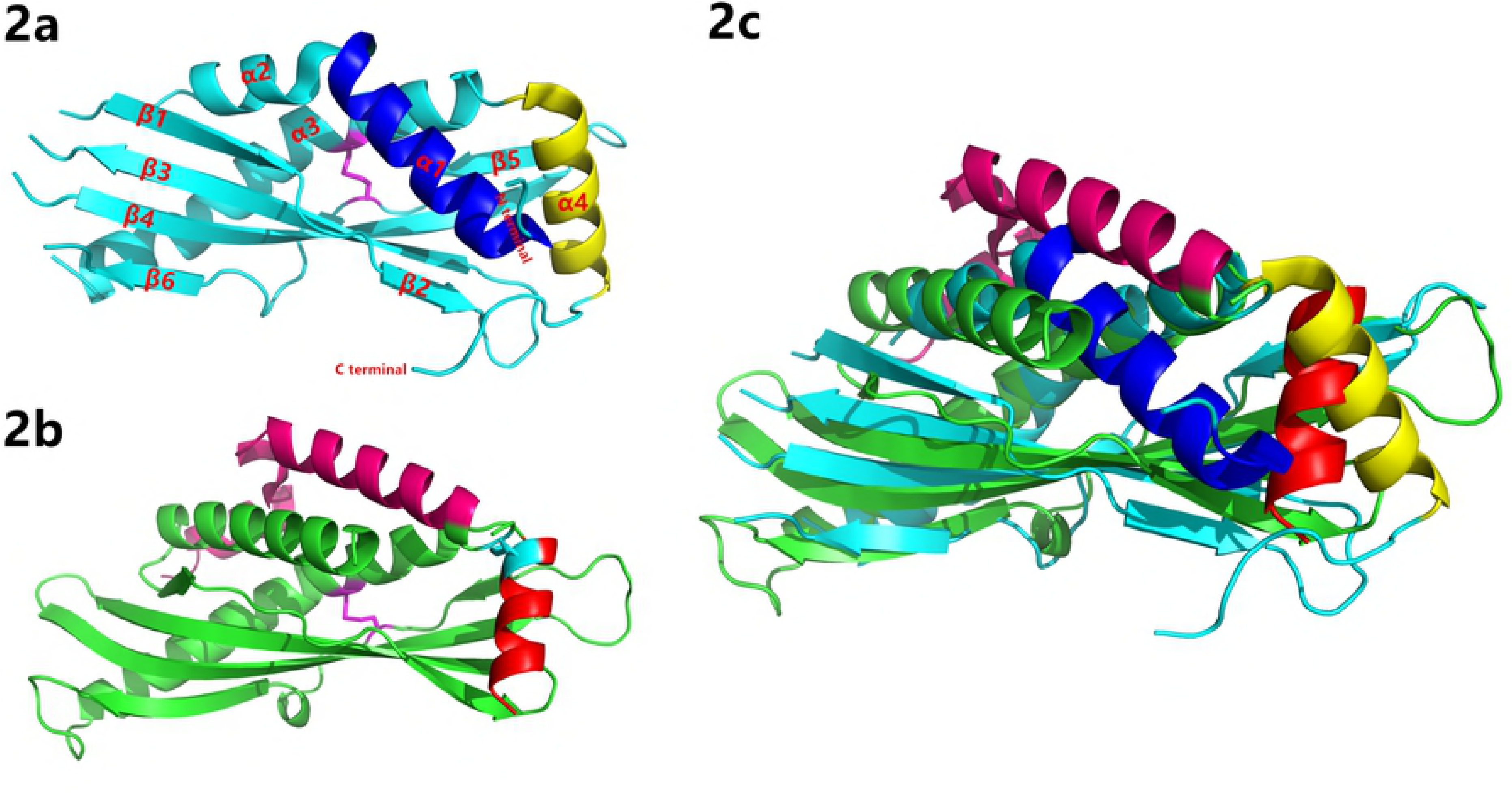
Structural comparison between BSP30b with hSPLUNC1. 2a, the structure of hSPLUNC1, with its disulfide bond shown in magenta. 2b, the structure of BSP30b. The conserved disulfide bond is shown in magenta and the second disulfide bond is shown is cyan. 2c, structural alignment of BSP30b to hSPLUNC1. BSP30b is shown in cyan and hSPLUNC1 is shown in green. The structural differences are marked with different colours as per 2a & b.

As SPLUNC1 and BPIFA1 are closely related, we take the comparison between BSP30b and SPLUNC1 as an example. There are two obvious disparities between BSP30b and SPLUNC1. The first difference results from the presence of a second disulfide bond in BSP30b (Fig. 2b). This second disulfide bond, formed between Cys72 and Cys229, connects the flexible loop just after helix α3 to the C-terminal helix α7 (Fig. 1b and 2b). Intriguingly the electron density for the disulphide bond is only clear in one molecule of the asymmetric unit and missing in the other. This may indicate that this disulfide bond is partially reduced under the crystallisation conditions.

The second obvious difference between the two structures is that both the N-terminus and C-terminus of SPLUNC1 are located at the same end of its long axis, whilst the termini of BSP30b are located at opposite ends of the long axis (Fig. 2c). That is to say that the first three N-terminal α helices in BSP30b have no structural equivalent in SPLUNC1. This structural difference influences the geometry of the opening of the ligand binding cavity. For BSP30b, the entrance to the ligand binding site is adjacent to the C-terminal helix α7 and is potentially influenced by the flexible loop between helices α3 and α4. Whereas for SPLUNC1, the entrance to the cavity is dominated by the long N-terminal helix (in blue in figure 2c).

The structure of BSP30b was also compared with the TULIP proteins whose structures have been deposited in the PDB. A sequence alignment of the corresponding PFam family (PF01273) (27) shows that the two cysteine residues that constitute the conserved disulphide bond are the only well conserved feature of the multiple sequence alignment. Hence the conservation of structure is the defining feature of this family. Generally, the secondary structures constituting the tubular channel are well conserved among all of the proteins. It is also clear that the C-terminal domains of BPI (1ewf), LBP (4m4d), and CETP (2obd) are well aligned but are different from E-syt 2 (4p42). This is because the C-terminal domain of E-syt 2 interacts with lipid bilayers, rather than lipid binding/transportation (28).

### Structure of BSP30b with oleic acid in the hydrophobic tunnel

Since the primary lipid sources for cattle are linolenic acid, linoleic acid, and oleic acid from pasture grass, we speculated that those fatty acids may be substrates for BSP30b. Thus, the binding of these three fatty acids were examined by co-crystallisation separately. Several structures were determined from crystals grown from mixtures of BSP30b and various fatty acids and fatty acid derivatives (e.g. methyl oleic ester). The only structure that revealed elongated stretches of electron density within the hydrophobic tunnel was the complex between BSP30b and oleic acid. The hydrophobic end of oleic acid is buried in the tunnel and its carboxyl group stretches out from the tunnel’s opening, forming hydrogen bonds with the carboxyl group of ASP112 and the carbonyl group of ILE113 (Fig. 3a and 3b). We suggest that the hydrophobic tail of oleic acid first “glides” into the internal cavity, driven by multiple hydrophobic interactions, while its carboxyl group stretches out and is stabilised by forming hydrogen bonds with the proximate amino acids. A similar hypothesis has been proposed for *Epiphyas postvittana* Takeout 1 protein (29), where both fatty acid ligands (including the head groups) “slide down” into the hydrophobic tunnel.

**Figure 3.**
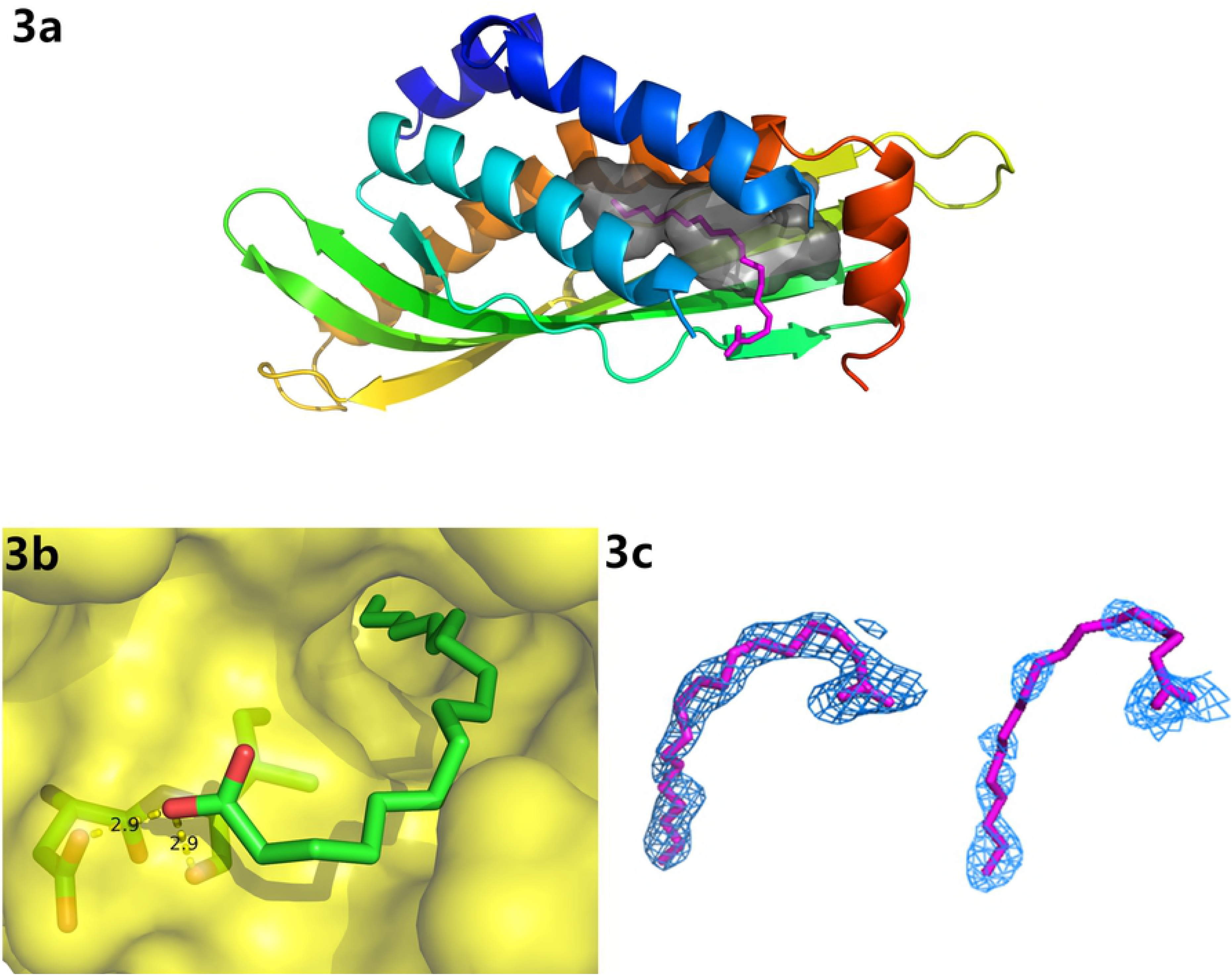
Structure of BSP30b with oleic acid. 3a Oleic acid is coloured in magenta and occupies the internal cavity, shown in grey. 3b, surface representation of the binding pocket with oleic acid. Hydrogen bonds between the carboxyl group of oleic acid and carboxyl group of ASP112, and backbone carbonyl group of ILE113 are shown as yellow dashed lines. 3c, electron density maps around both oleic acid molecules in the asymmetric unit. The final σa-weighted 2mFo-DFc map is contoured at 0.8 σ (blue). All structural illustrations were prepared with PyMOL.

We also observe that the overall structure of BSP30b and its internal cavity remain almost completely unchanged upon binding of the lipid ligand, suggesting that the internal cavity of BSP30b also forms a rigid scaffold for its lipid ligands. Only the helix α3 moves slightly away. It is noteworthy that the electron density of oleic acid is clear in one molecule of the asymmetric unit but incomplete for oleic acid in the other molecule (Fig. 3c). This may suggest that oleic acid is not the preferred ligand of BSP30b. A refinement of the occupancy for the binding site of the predominant ligand showed 80% occupancy for this molecule.

### GFP_BSP30b is an emulsifier with olive oil

Twenty minutes after mixing GFP_BSP30b with olive oil, two layers were formed. The top layer was thin and cloudy whereas the bottom layer was slightly cloudy. When the bottom layer was viewed under blue light by fluorescent microscopy, an emulsion was seen with GFP_BSP30b decorating the surface of each small oil droplet, indicated by its green fluorescence (Fig. 4a). The top thin layer also has a similar composition but the small discreet droplets packed against one another. In decorating the surface of oil droplets, BSP30b may stabilise the small oil droplets in solution and thus, act as an emulsifier. This observation is consistent with the original finding by Wheeler and colleagues (8) that BSP proteins are protective for bloat in cattle.

**Figure 4.**
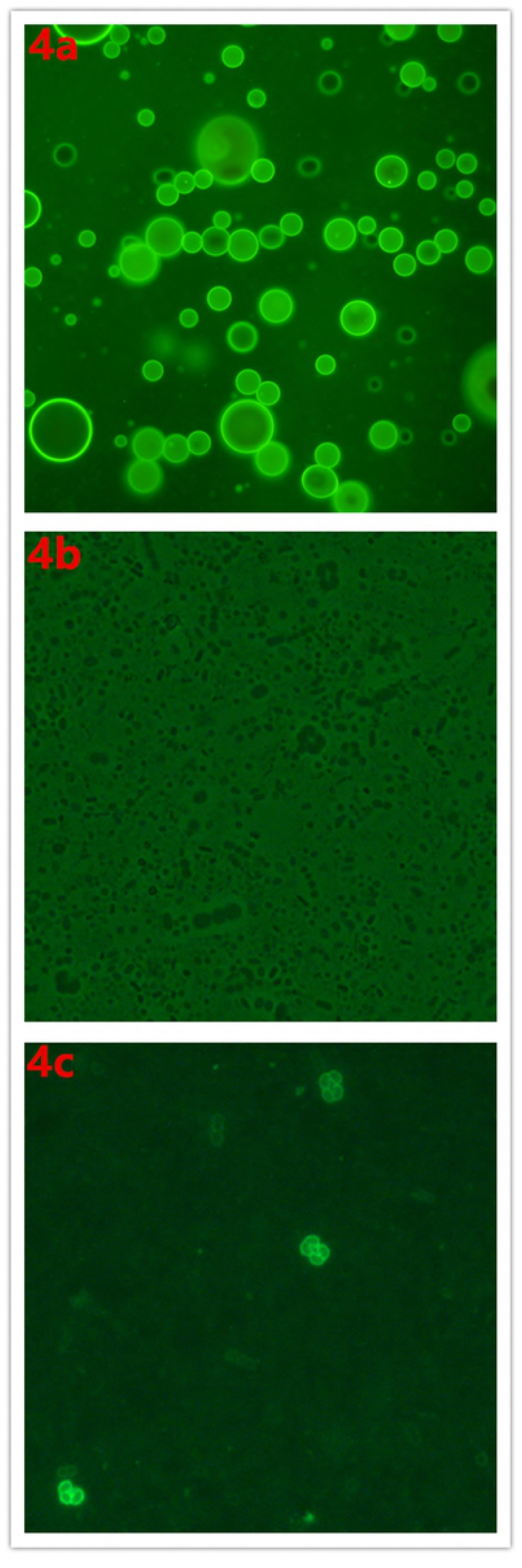
Fluorescent microscopy of GFP_BSP30b coating olive oil/rumen bacteria. 4a, emulsion formed after mixing of GFP_BSP30b with olive oil. BSP30b decorates the surface of each droplet, indicated by green fluorescence from its fused GFP under UV illumination. 4b and 4c, comparison of the view of rumen bacteria mixed with GFP_BSP30b under bright field (4b) and UV illumination (4c).

### GFP_BSP30b binding to the surface of specific rumen bacterial strains

A surprising result to us is the observation that the mixture of GFP_BSP30b with ruminant bacteria observed under UV illumination shows that BSP30b binds to just a small number of rumen bacterial cells. BSP30b showed strong binding to a short and curved rod-like bacterial strain, which is clustered as triad or tetrad (Fig. 4c). Whereas BSP30b does not bind to the majority of bacteria in the field of view. We observed that BSP30b can also bind to a few cocci-319 shaped cells but with weaker fluorescence suggesting a weaker equilibrium constant for binding to these bacteria. We suggest that the binding specificity of BSP30b to certain bacterial surface lipids may result in its binding selectivity to a small subset of rumen bacterial strains and characterising these strains is the subject of ongoing work in our laboratory.

## Discussion

Currently, the bovine PLUNC family has 13 members including BSP30a-d that have likely arisen through recent gene duplication (13). Although BSP30 gene expression has been well-studied at both the mRNA level and the protein level, their biological functions are poorly understood. The lack of structural information or knowledge about the lipid substrates for the cattle PLUNC family members impedes our understanding of their function.

Here, we report the first crystal structures of BSP30b at 2.0 Å resolution. We have shown that the TULIP superfamily architecture is maintained with the characteristic internal hydrophobic channel formed by an α/β wrap fold. Structural comparison to the closest related structures in the PDB (human SPLUNC1 and mouse BPIFA1) has revealed both conserved and unique features that might be important for function. The unique disulfide bond present in BSP30b but not in other PLUNC proteins may play a role in ligand binding as it links the C-terminal helix with the protein body and the C-terminal helix lies adjacent to the ligand tunnel. Further evidence for this includes the fact that a similar disulfide bond is also found in the structure of Takeout 1 of *Epiphyas postvittana* and JHBP of silkworm, which are also members of the TULIP superfamily. Their disulfide bonds are reported to have a critical role in ligand binding rather than maintaining the stability of the bulk structure (29, 30).

Compared to human SPLUNC1, the internal cavity opening of BSP30b is significantly larger. This results from the reorganisation of the termini of the protein such that the C-terminus of BSP30b lies at the entrance to the ligand binding tunnel. For SPLUNC1, its N-terminal helix α1 and C-terminal helix α4, together with helix α3, form the predicted ligand binding site of SPLUNC1 and it is hypothesized that helix α1 moves away to “open-up” the binding site upon lipid binding (26). From our BSP30b-oleic acid structure, it is obvious that, unlike the changes proposed for SPLUNC1, no major conformational change is needed for lipid binding. Only helix α3 has moved away slightly from the core structure upon oleic acid binding. Ligand binding may be related to oxidation/reduction of the disulphide bond that links the C-terminal helix in a similar manner to the takeout proteins.

This is also the first time that a PLUNC family protein has been crystallized with its ligand, although the electron density for the ligand suggests ∼80% occupancy. We have tried to crystalize BSP30b with linolenic acid and linoleic acid, but in the structural data no electron density was found for either of these fatty acids. Linolenic acid and linoleic acid have more rigid structures compared to oleic acid as they have three and two double bonds on their carbon backbones, respectively. In contrast, oleic acid has only one double bond, which gives it more flexibility to occupy the relatively narrow hydrophobic channel.

Proteins in the PLUNC family have been shown to have potent surfactant properties at water/liquid interfaces (31, 32). We demonstrated that BSP30b also behaves like an emulsifier, dispersing olive oil into small droplets in aqueous solution. This may be relevant to the original observation that high BSP30 expressing animals had lower susceptibility to bloat (which is caused by uncontrolled foaming in the rumen).

We find it intriguing that BSP30b only binds to a small subset of bacterial cells within the rumen. “Bacteria coating” by proteins from the PLUNC group has been reported before (33, 34). The binding of murine PSP to *E. coli*, *S. mutans, A. actinomycetemcomitans*, and *L. monocytogenes* has been demonstrated to be protein-protein interactions dependent on the presence of Zn^2+^. However, we suggest that BSP30b binds to specific lipids on the bacterial surface based on the crystal structure and oleic acid binding. The “bacterial cell coating” by human SPLUNC1 on *P. aeruginosa*-pMF230 has been investigated using confocal microscopy, and the protein is proposed to display a bacteriostatic property preventing bacterial growth. Other studies also report that anti-biofilm properties of human SPLUNC1 may due to its role as a surfactant (35). In light of our structure and functional experiments with BSP30b we suggest that this abundant salivary protein could have important roles in the rumen via its binding specificity to certain ruminal bacteria species and identifying the specific bacterial species and the nature of the interaction is the subject of ongoing work.

## Acknowledgements

We would like to thank DairyNZ (Hamilton, New Zealand) for their funding. This research was undertaken in part using the MX2 beamline at the Australian Synchrotron, part of ANSTO, and made use of the Australian Cancer Research Foundation (ACRF) detector.

## Abbreviations

BSP30b: bovine salivary protein 30b
PLUNC: palate, lung, and nasal epithelium carcinoma-associated
BPI: bactericidal/permeability-increasing
TULIP: tubular lipid-binding

## References

1. Canny G, Levy O. Bactericidal/permeability-increasing protein (BPI) and BPI homologs at mucosal sites. Trends Immunol. 2008 Nov;29(11): 541–7. PubMed PMID: 18838299.

2. LeClair EE. Four reasons to consider a novel class of innate immune molecules in the oral epithelium. J Dent Res. 2003 Dec;82(12): 944–50. PubMed PMID: 14630892.

3. Ghafouri B, Stahlbom B, Tagesson C, Lindahl M. Newly identified proteins in human nasal lavage fluid from non-smokers and smokers using two-dimensional gel electrophoresis and peptide mass fingerprinting. Proteomics. 2002 Jan;2(1): 112–20. PubMed PMID: 11788998.

4. Ghafouri B, Kihlstrom E, Tagesson C, Lindahl M. PLUNC in human nasal lavage fluid: Multiple isoforms that bind to lipopolysaccharide. Biochim et Biophys Acta. 2004 Jun 1;1699(1-2):57–63. PubMed PMID: 15158712.

5. Bartlett JA, Hicks BJ, Schlomann JM, Ramachandran S, Nauseef WM, McCray PB, Jr. PLUNC is a secreted product of neutrophil granules. J Leukocyte Biol. 2008 May;83(5): 1201–6. PubMed PMID: 18245229.

6. Kopec KO, Alva V, Lupas AN. Bioinformatics of the TULIP domain superfamily. Biochem Soc Trans. 2011 Aug;39(4): 1033–8. PubMed PMID: 21787343.

7. Alva V, Lupas AN. The TULIP superfamily of eukaryotic lipid-binding proteins as a mediator of lipid sensing and transport. Biochim et Biophys Acta. 2016 Aug;1861(8 Pt B):913–23. PubMed PMID: 26825693.

8. Rajan GH, Morris CA, Carruthers VR, Wilkins RJ, Wheeler TT. The relative abundance of a salivary protein, BSP30, is correlated with susceptibility to bloat in cattle herds selected for high or low bloat susceptibility. Anim Genet. 1996 Dec;27(6): 407–14. PubMed PMID: WOS:A1996WD14700003.

9. Clarke RT, Reid CS. Foamy bloat of cattle. A review. J Dairy Sci. 1974 Jul;57(7): 753–85. PubMed PMID: 4601637.

10. Bingle CD, Seal RL, Craven CJ. Systematic nomenclature for the PLUNC/PSP/BSP30/SMGB proteins as a subfamily of the BPI fold-containing superfamily. Biochemi Soc Trans. 2011 Aug;39(4): 977–83. PubMed PMID: 21787333. Pubmed Central PMCID: 3196848.

11. Wheeler TT, Haigh BJ, McCracken JY, Wilkins RJ, Morris CA, Grigor MR. The BSP30 salivary proteins from cattle, LUNX/PLUNC and von Ebner’s minor salivary gland protein are members of the PSP/LBP superfamily of proteins. Biochim et Biophys Acta. 2002 Dec 12;1579(2-3):92–100. PubMed PMID: 12427544.

12. Wheeler TT, Hood KA, Maqbool NJ, McEwan JC, Bingle CD, Zhao SY. Expansion of the bactericidal/permeability increasing-like (BPI-like) protein locus in cattle. BMC Genomics. 2007 Mar 15;8. PubMed PMID: WOS:000245454600002.

13. Haigh B, Hood K, Broadhurst M, Medele S, Callaghan M, Smolenski G, et al. The bovine salivary proteins BSP30a and BSP30b are independently expressed BPI-like proteins with anti-Pseudomonas activity. Mol Immunol. 2008 Apr;45(7): 1944–51. PubMed PMID: WOS:000254260500013.

14. Wheeler TT, Hood K, Oden K, McCracken J, Morris CA. Bovine parotid secretory protein: Structure, expression and relatedness to other BPI (bactericidal/permeability-increasing protein)-like proteins. Biochem Soc Trans. 2003 Aug;31(Pt 4):781–4. PubMed PMID: 12887305.

15. Wheeler TT, Haigh BJ, Broadhurst MK, Hood KA, Maqbool NJ. The BPI-like/PLUNC family proteins in cattle. Biochem Soc Trans. 2011 Aug;39:1006–11. PubMed PMID: WOS:000293814800028.

16. Leslie AGW, Powell HR. Processing diffraction data with MOSFLM. Nato Sci Ser Ii-Math. 2007;245:41-+. PubMed PMID: WOS:000248330600004.

17. Evans PR, Murshudov GN. How good are my data and what is the resolution? Acta Crystallogr D. 2013 Jul;69:1204–14. PubMed PMID: WOS:000320712800003.

18. Collaborative Computational Project N. The CCP4 suite: Programs for protein crystallography. Acta Crystallogra D. 1994 Sep 1;50(Pt 5):760–3. PubMed PMID: 15299374.

19. Terwilliger TC, Adams PD, Read RJ, Mccoy AJ, Moriarty NW, Grosse-Kunstleve RW, et al. Decision-making in structure solution using Bayesian estimates of map quality: the PHENIX AutoSol wizard. Acta Crystallogr D. 2009 Jun;65:582–601. PubMed PMID: WOS:000266216900009.

20. Terwilliger TC, Grosse-Kunstleve RW, Afonine PV, Moriarty NW, Zwart PH, Hung LW, et al. Iterative model building, structure refinement and density modification with the PHENIX AutoBuild wizard. Acta Crystallogr D. 2008 Jan;64:61–9. PubMed PMID: WOS:000251357200008.

21. Adams PD, Afonine PV, Bunkoczi G, Chen VB, Echols N, Headd JJ, et al. The Phenix software for automated determination of macromolecular structures. Methods. 2011 Sep;55(1): 94–106. PubMed PMID: WOS:000296038800012.

22. Afonine PV, Grosse-Kunstleve RW, Adams PD. A robust bulk-solvent correction and anisotropic scaling procedure. Acta Crystallogr D. 2005 Jul;61:850–5. PubMed PMID: WOS:000230113000002.

23. Emsley P, Cowtan K. Coot: model-building tools for molecular graphics. Acta Crystallogr D. 2004 Dec;60:2126–32. PubMed PMID: WOS:000225360500002.

24. McPhillips TM, McPhillips SE, Chiu HJ, Cohen AE, Deacon AM, Ellis PJ, et al. Blu-Ice and the Distributed Control System: software for data acquisition and instrument control at macromolecular crystallography beamlines. J Synch Radiation. 2002 Nov 1;9(Pt 6):401–6. PubMed PMID: 12409628.

25. Mccoy AJ, Grosse-Kunstleve RW, Adams PD, Winn MD, Storoni LC, Read RJ. Phaser crystallographic software. J Appl Crystallogr. 2007 Aug;40:658–74. PubMed PMID: WOS:000248077500003.

26. Ning FK, Wang C, Berry KZ, Kandasamy P, Liu HL, Murphy RC, et al. Structural characterization of the pulmonary innate immune protein SPLUNC1 and identification of lipid ligands. FASEB J. 2014 Dec;28(12): 5349–60. PubMed PMID: WOS:000345894500028.

27. Finn RD, Coggill P, Eberhardt RY, Eddy SR, Mistry J, Mitchell AL, et al. The Pfam protein families database: Towards a more sustainable future. Nucleic Acids Res. 2016 Jan 4;44(D1):D279–85. PubMed PMID: 26673716.

28. Schauder CM, Wu X, Saheki Y, Narayanaswamy P, Torta F, Wenk MR, et al. Structure of a lipid-bound extended synaptotagmin indicates a role in lipid transfer. Nature. 2014 Jun 26;510(7506): 552–5. PubMed PMID: 24847877.

29. Hamiaux C, Basten L, Greenwood DR, Baker EN, Newcomb RD. Ligand promiscuity within the internal cavity of *Epiphyas postvittana* Takeout 1 protein. J Struct Biol. 2013 Jun;182(3): 259–63. PubMed PMID: 23563188.

30. Suzuki R, Fujimoto Z, Shiotsuki T, Tsuchiya W, Momma M, Tase A, et al. Structural mechanism of JH delivery in hemolymph by JHBP of silkworm, *Bombyx mori*. Sci Reports. 2011;1:133. PubMed PMID: 22355650.

31. Bartlett JA, Gakhar L, Penterman J, Singh PK, Mallampalli RK, Porter E, et al. PLUNC: a multifunctional surfactant of the airways. Biochem Soc Trans. 2011 Aug;39(4): 1012–6. PubMed PMID: 21787339.

32. McDonald RE, Fleming RI, Beeley JG, Bovell DL, Lu JR, Zhao X, et al. Latherin: a surfactant protein of horse sweat and saliva. PLoS One. 2009 May 29;4(5):e5726. PubMed PMID: 19478940.

33. Robinson CP, Bounous DI, Alford CE, Nguyen KH, Nanni JM, Peck AB, et al. PSP expression in murine lacrimal glands and function as a bacteria binding protein in exocrine secretions. Amer J Physiol. 1997 Apr;272(4 Pt 1):G863–71. PubMed PMID: 9142919.

34. Sayeed S, Nistico L, St Croix C, Di YP. Multifunctional role of human SPLUNC1 in *Pseudomonas aeruginosa* infection. Infect Immun. 2013 Jan;81(1): 285–91. PubMed PMID: 23132494.

35. Gakhar L, Bartlett JA, Penterman J, Mizrachi D, Singh PK, Mallampalli RK, et al. PLUNC is a novel airway surfactant protein with anti-biofilm activity. PLoS One. 2010 Feb 9;5(2):e9098. PubMed PMID: 20161732.

